# Pentamidine inhibition of streptopain attenuates *Streptococcus pyogenes* virulence

**DOI:** 10.1101/2025.03.12.642885

**Authors:** Keya Trivedi, Christopher N. LaRock

## Abstract

The obligate human pathogen Group A *Streptococcus* (GAS; *Streptococcus pyogenes*) carries high morbidity and mortality, primarily in impoverished or resource-poor regions. The failure rate of monotherapy with conventional antibiotics is high, and invasive infections by this bacterium frequently require extensive supportive care and surgical intervention. Thus, it is important to find new compounds with adjunctive therapeutic benefits. The conserved secreted protease streptopain (Streptococcal pyrogenic exotoxin B; SpeB) directly contributes to disease pathogenesis by inducing pathological inflammation, degrading tissue, and promoting the evasion of antimicrobial host defense proteins. This study screened 400 diverse off-patent drug and drug-like compounds for inhibitors of streptopain proteolysis. Lead compounds were tested for activity at lower concentrations and anti-virulence activities during *in vitro* infection. Significant inhibition of streptopain was seen for pentamidine, an anti-protozoal drug approved for the treatment of pneumocystis pneumonia, leishmaniasis, and trypanosomiasis. Streptopain inhibition rendered GAS susceptible to killing by human innate immune cells. These studies identify unexploited molecules as new starting points for drug discovery and a potential for repurposing existing drugs for the treatment of infections by GAS.

**IMPORTANCE:** *Streptococcus pyogenes* (group A strep; GAS) is a common cause of severe invasive infections. Repeated infections can trigger autoimmune diseases such as acute rheumatic fever and rheumatic heart disease. This study examines how targeting a specific, highly conserved virulence factor of the secreted cysteine protease streptopain, can sensitize a serious pathogen to killing by the immune system. Manipulating the host-pathogen interaction, rather than attempting to directly kill a microbe, is a promising therapeutic strategy. Notably, its benefits include limiting off-target effects on the microbiota and potentially limiting the development and spread of resistance. Streptopain inhibitors, including the antifungal and antiparasitic drug pentamidine as identified in this work, may therefore be useful in the treatment of *S. pyogenes* infection.

## INTRODUCTION

Group A *Streptococcus* (GAS, *Streptococcus pyogenes*) is a top cause of infectious mortality and is responsible for over half a million annual deaths worldwide (1). Humans are colonized throughout childhood, leading to bouts of pharyngitis or pyoderma, but often without overt symptoms of disease (2). Rheumatic heart disease can follow repeated infections, while any infection can become more invasive, leading to cellulitis, necrotizing fasciitis, scarlet fever, or sepsis. GAS remains sensitive to penicillin antibiotics, which are a mainstay of pharyngitis (“strep throat”). However, this monotherapy is typically insufficient during invasive infections, which is highly pro-inflammatory (3) and can require adjunctive antibiotics, surgical removal of infected tissue, and extensive supportive care (4). The case fatality of these severe infections is high in the United States and worse in resource-limited environments where infection is common.

The secreted cysteine protease SpeB is a primary virulence factor of GAS and is an attractive drug target for several reasons (5). Among these, SpeB is unique to GAS, but highly conserved within the species, is abundant during infection (6–8), and correlates with the severity of disease in humans and in model infections (9–13). *In vitro* and *in vivo* experiments show that SpeB degrades tissue (14, 15), inactivates immune antimicrobials (13, 16–24), and activates pathological proinflammatory pathways through direct and indirect mechanisms (25–30). These activities cumulatively contribute to both GAS growth and injury to the host (31). Consequently, SpeB promotes pathogenesis in skin (32–34) and upper respiratory (26, 35) infection models in mice.

The aim of this study was to identify small molecule inhibitors of SpeB that would have therapeutic potential for treating infections by GAS. The catalytic cysteine of SpeB, like most other cysteine proteases, is highly reactive and can be inhibited by most common inhibitors of this family (5, 36). However, broad-spectrum inhibitors would be unsuitable as anti-infectives due to targeting of mammalian proteases leading to toxicity and immunosuppressive effects (37). Furthermore, in addition to looking for strong activity against SpeB, we also considered potential accessibility to the populations where it would be most needed. To these ends, we screened a diverse library of off-patent drug and drug-like compounds that was developed by the Medicines for Malaria Venture for the treatment of poverty-related diseases. All compounds of the library have been screened for toxicity (38), a large fraction of them have been tested *in vivo*, and >10% are approved for use in humans for at least one indication. Most potential drugs fail in pre-clinical or clinical studies; repurposing of existing pharmaceuticals is an avenue to potentially bypass this bottleneck inexpensively, quickly, and safely. We define several lead compounds with therapeutic potential.

## MATERIALS AND METHODS

### Bacterial strains

GAS 5448, *S. aureus* F-182, and their growth are previously described (39–41). Briefly, bacteria were routinely grown in Todd-Hewitt Yeast (THY; Difco) or on THY-agar at 37°C and 5% CO_2_. Bacteria from overnight cultures were first washed and diluted in PBS in preparation for infection experiments. Native SpeB was purified from culture supernatants by ammonium sulfate (Sigma) precipitation followed by cleanup by size-exclusion chromatography (ÄKTA FPLC, PE) and Centricon filtration (EMD Millipore), as previously described (24).

### Chemical library

The Pathogen Box library compound was kindly provided by Medicines for Malaria Venture (MMV, Switzerland) dissolved in DMSO and arrayed in 96-well format. Stocks were duplicated in DMSO and stored at-80°C, and dilutions were made fresh for each experiment. Test compounds in this library were selected as having a minimum of 5-fold selectivity over mammalian cells for either *Plasmodium, Mycobacterium tuberculosis*, or kinetoplastid protists. 26 additional drugs, including conventional antibiotics (eg. doxycycline, rifampin, streptomycin, linezolid), anti-infectives against non-bacterial pathogens (eg. praziquantel, nifurtimox, miltefosine), and other common drugs (eg. fluoxetine, auranofin) were included as an additional reference set. The lead hit pentamidine was validated with additional high purity compounded purchased from commercial supplier (Sigma).

### Molecular docking

The interaction of mature SpeB sequence of PDB:2uzj, and the chemical structures of each antibiotic from PubChem was examined using the Chai-1 Model (42). Modeling was agnostic with no input restraints specified, and the best scoring model was selected for analysis in ChimeraX. Each molecule was oriented to the same position using Matchmaker to displaying antibiotic occupancy in the substrate cleft of the enzyme, and red and blue used to indicate computed surface electrostatics, as previously described (43).

### SpeB inhibitor screen

In the initial screen, SpeB activity was measured in the presence of test compound in individual wells using the substrate azocasein (Sigma) essentially as previously described (32). Briefly, test compounds were added to achieve a 100 μM final concentration with 10 nM SpeB and 2% (w/v) casein labeled with azo dye (Sigma), all in PBS pH 7.4 with 2 mM dithiothreitol (Sigma). Reactions were incubated 18 h at 37°C. Inhibition was observed if the opacity of the solution remained comparable to controls to which no SpeB was added.

### SpeB inhibitor validation

Kinetics of SpeB inhibition by compounds passing the first screen were measured using the internally-quenched peptide sub103, Mca-IFFDTWK-Dnp (CPC Scientific), as previously (32). In triplicate, dilutions of peptides were incubated in PBS with 5 mM dithiothreitol and 0.01% Tween 20 with 10 nM SpeB and dilutions of test compound. The reaction was incubated at 37°C and continuously monitored with fluorophore excitation at 323 nm and emission at 398 nm on a Nivo plate reader (PerkinElmer).

### Neutrophil infection assays

Whole human blood was collected from healthy adult donors with informed consent and approval from the Institutional Review Board at Emory University. Blood was collected into heparinized Vacutainer tubes, and primary neutrophils were isolated using PolymorphPrep (Axis-Shield). Neutrophils were diluted to 10^5^ cells/mL in RPMI containing 10% FBS with no antibiotic, infected with 1 × 10^6^ CFU of GAS for 60 minutes, then plated on THY agar, essentially as previously (44). CFU were enumerated after overnight incubation of THY agar plates at 37°C.

### Statistical analysis

Statistical analyses were performed using Prism 10 (GraphPad). Values are expressed as means ± standard error unless otherwise specified. Differences were determined using the Mann-Whitney U test (paired) or the Kruskal-Wallis test with Dunn’s post-analysis (multiple groups) unless otherwise specified.

## RESULTS

### Initial screen

The complex regulation of streptopain/SpeB (45–48) can lead to identification of compounds whose impact on SpeB is indirect if tested in culture with live bacteria (49–51). Therefore, we conducted a screen using purified SpeB to focus only on those compounds that directly impact on the enzyme’s ability to hydrolyze substrates. Azo dye-labeled casein was incubated with purified SpeB in the presence of 400 compounds, each at 100 μM, and conversion of the chromogenic substrate measured after 18 h incubation at 37°C. 9 test compounds inhibited SpeB activity >50% across 2 independent runs (**Figure 1A**), for a hit rate of ∼2% (**Figure 1B**). No compound significantly increased apparent proteolysis of the azocasein substrate by SpeB. Compounds were selected for this library based on having known bioactivity toward one or more pathogens, possibly contributing to this enriched hit rate relative to similar screens (52).

**FIG 1.**
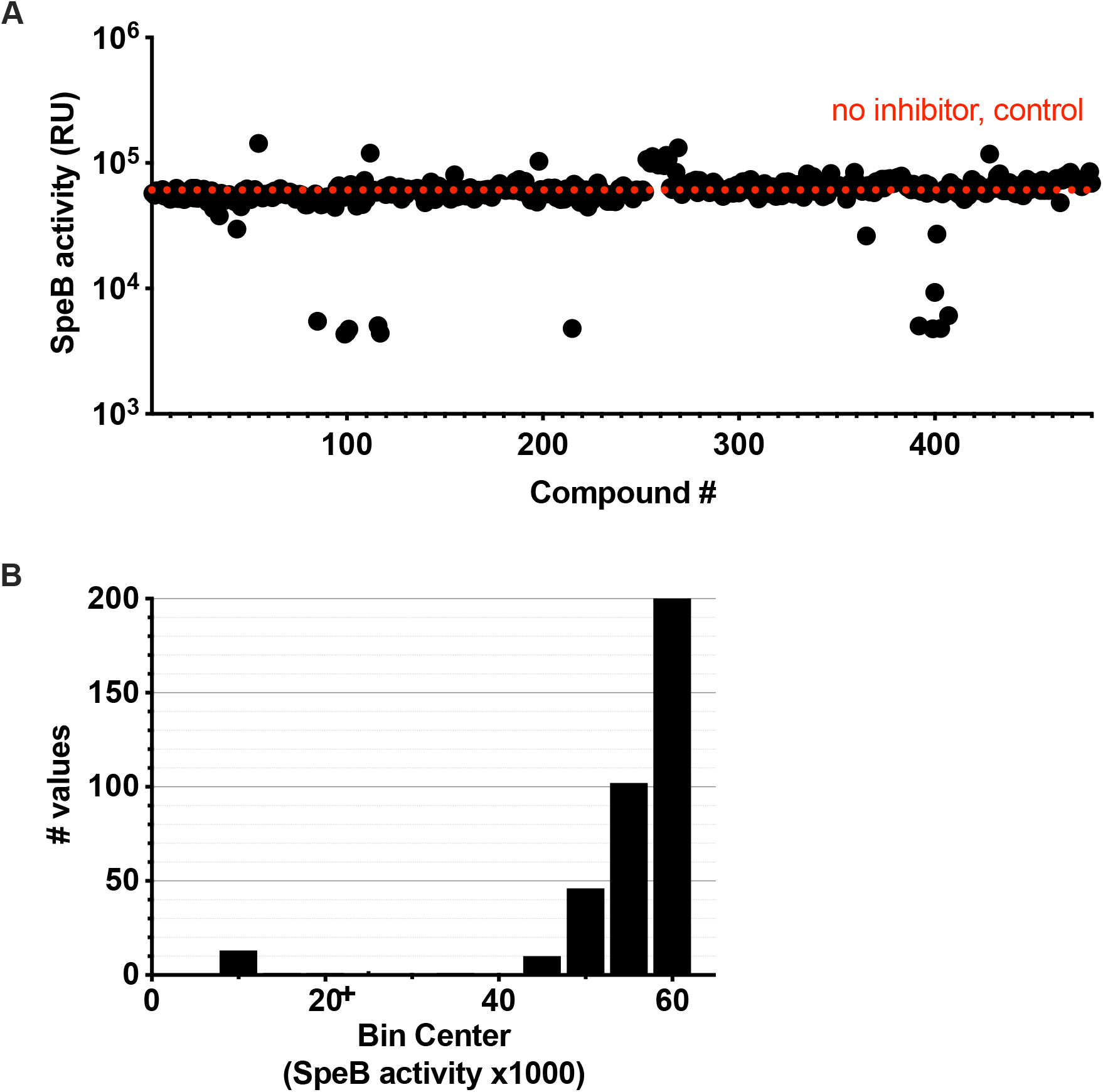
Graphical representation of results from the primary screen for each of the 400 Pathogen compounds. (A) Each individual compound was tested at a concentration of 100 μM for hydrolysis of the substrate azo-casein. (B) Distribution of results.

### Inhibition of SpeB by library compounds

Conversion of azocasein is a measurement of cumulative substrate degradation over a prolonged period of time (32). To more sensitively monitor the kinetics of proteolysis, we used the internally-quenched Förster resonance energy transfer (FRET) peptide substrate sub103, which is highly selective and only contains a single cleavage site for SpeB (32). Under these more stringent conditions, all compounds retained inhibitory activity (**Figure 2**), but not to sub-micromolar concentrations, a typical target for antiinfective drugs (37, 38). Nonetheless, these can be used to inform lead compounds for further chemical refinement.

**FIG 2.**
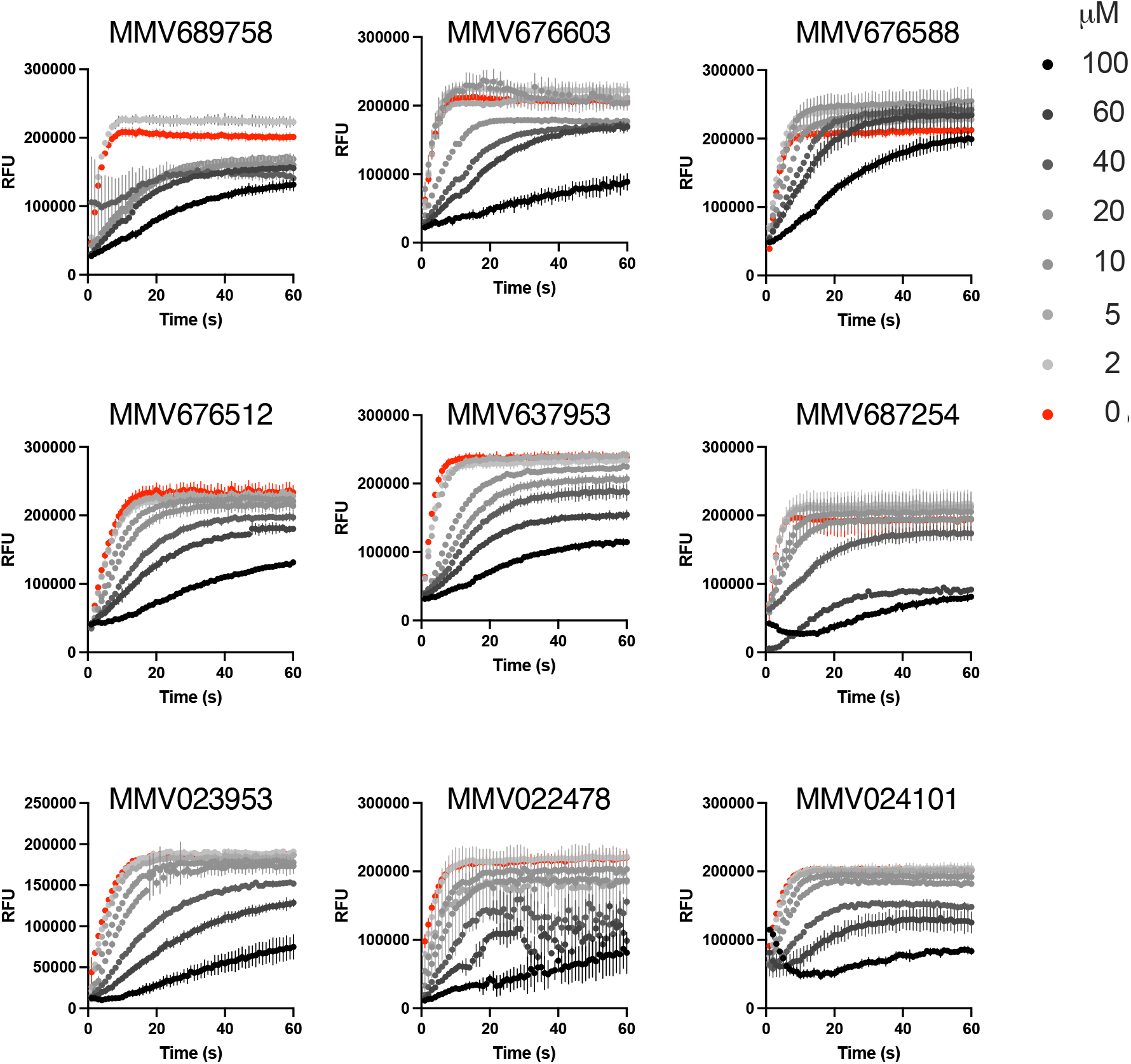
Activity of candidate compounds against SpeB. Cleavage of sub103 peptide after 30 min incubation with purified SpeB with each compound at the concentration indicated. N=4, error bars represent standard deviations.

### Structural similarity analysis

To determine whether there was any commonality between SpeB inhibitors, with the expectation that compounds with similar chemical structures could be used to identify additional compounds for testing, used molecular structure prediction modeling to examine their predicted interaction with SpeB. By unguided analysis, each compound was predicted to occupy the same region in the substrate pocket of SpeB (**Figure 3**). There was limited structural homology between compounds, but this modality was consistent with the kinetic analysis (**Figure 2**) that these compounds are competitive inhibitors for substrate binding.

**FIG 3.**
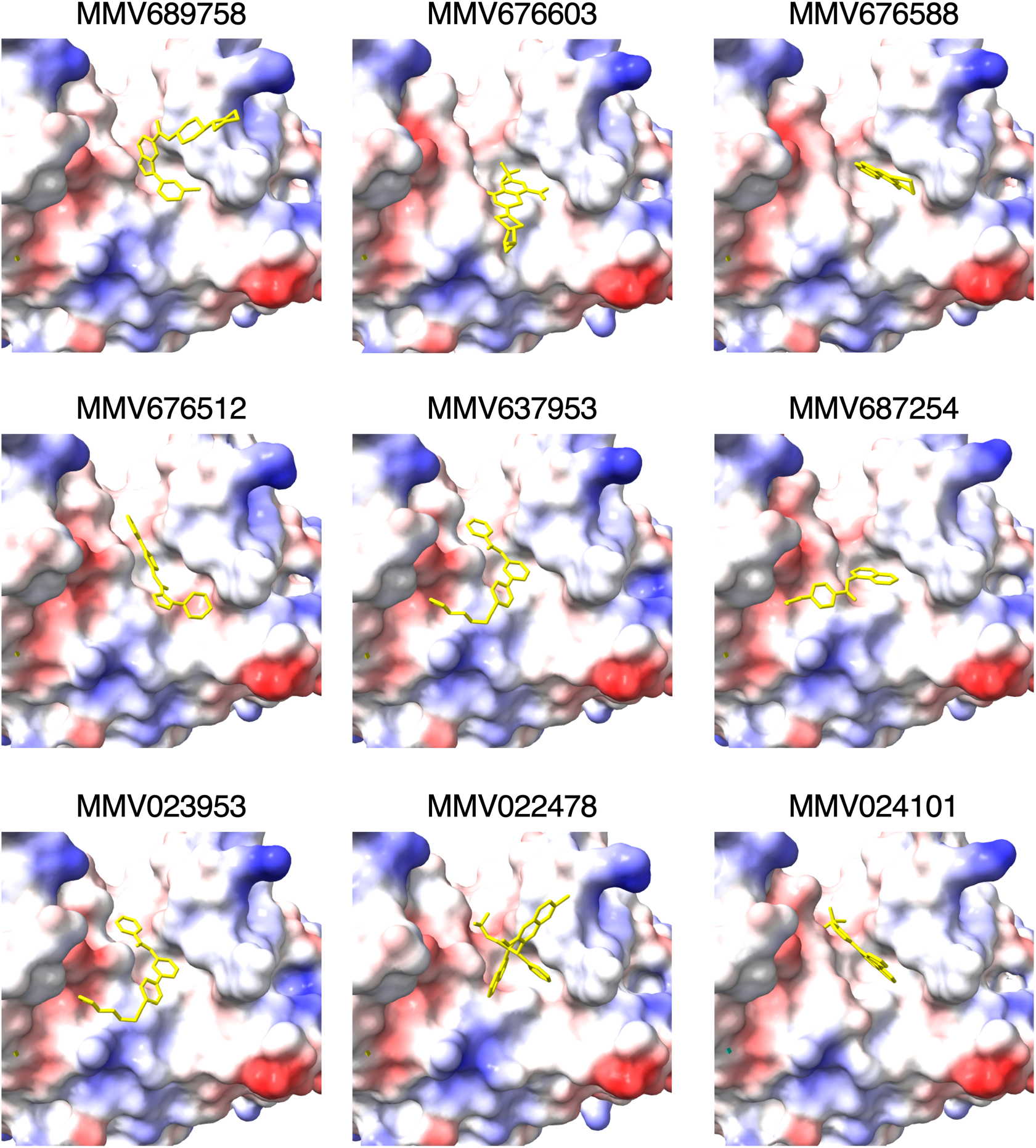
SpeB-inhibitor interactions. Models of the substrate pocket of SpeB and each indicated compound (yellow). Protein surface electrostatics are colored red and blue for negative and positive charges, respectively, and white color represents neutral residues.

### Inhibition of SpeB by pentamidine

Since this library was targeted for neglected tropical diseases, it additionally contained pentamidine as a positive control. While it is effective against several eukaryotic pathogens, efficacy against bacteria and their toxins, and SpeB of GAS in particular, has not been established. Pentamidine was predicted to occupy the catalytic site of SpeB similarly to the established inhibitor E64, in its cocrystal structure (53) and by prediction (**Figure 4A, 4B, 4C**). Further testing pentamidine’s efficacy at SpeB inhibition with titrations of drug, we find an IC_50_ of 9.671e-09 (**Figure 4D**). This was suggestive that pentamidine could be effective at inhibiting SpeB.

**FIG 4.**
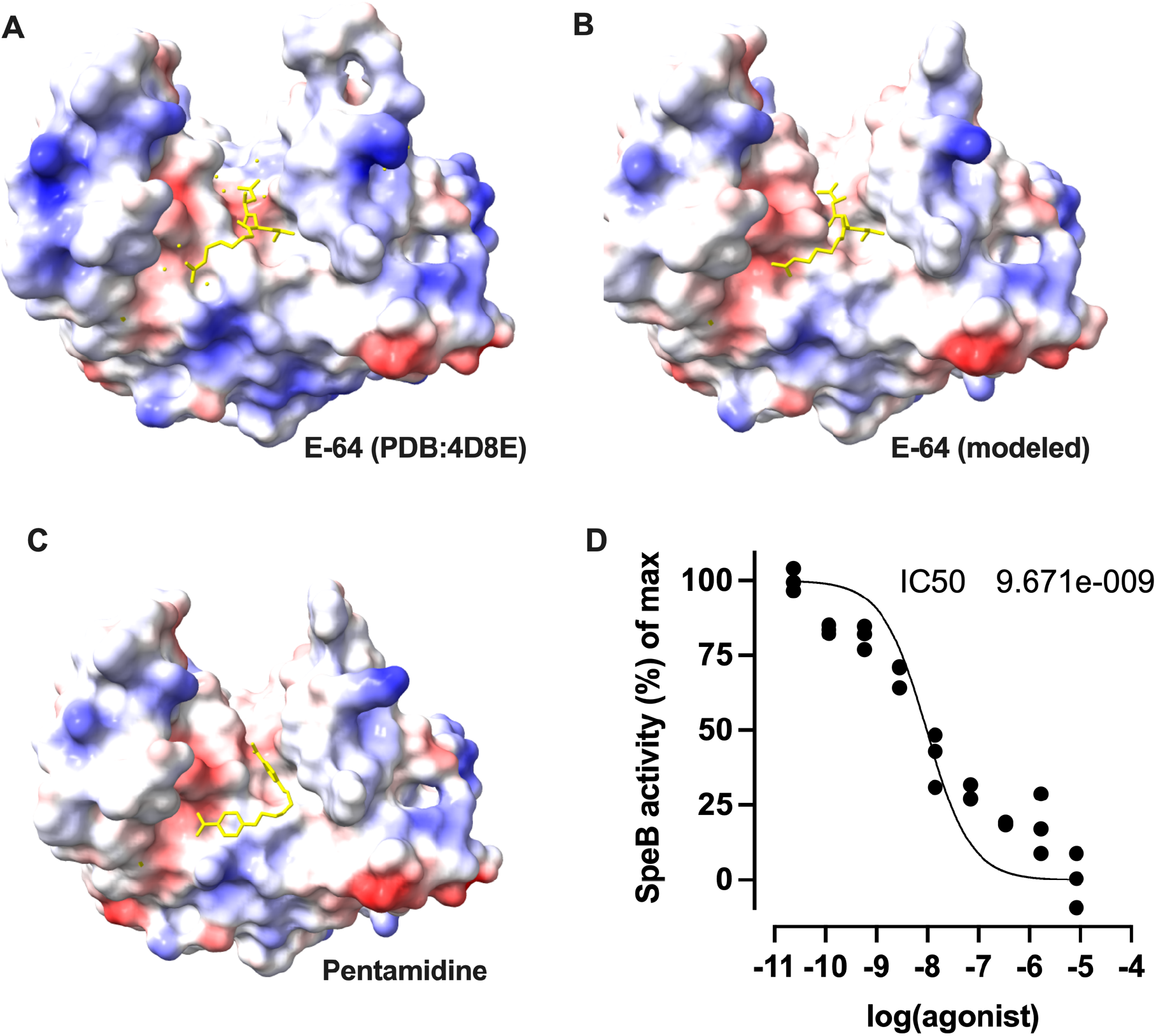
Pentamidine inhibits proteolysis by SpeB. (A-C) Models of the substrate pocket of SpeB and each indicated compound (yellow). Protein surface electrostatics are colored red and blue for negative and positive charges, respectively, and white color represents neutral residues. (D) Cleavage of sub103 peptide after 30 min incubation with purified SpeB was plotted against a range of pentamidine concentrations. N=4, error bars represent standard deviations.

### *In vitro* activity of pentamidine against *S. pyogenes*

SpeB is important for GAS resistance to innate immune antimicrobials, including those produced by neutrophils (46, 54). To examine whether SpeB inhibition would be useful for sensitizing GAS to neutrophil killing, we modeled this important host-pathogen interaction *in vivo*. 10 μM had no impact on the growth of GAS (**Figure 5A**) or *Staphylococcus aureus*, a bacteria included as a control for species-specific effects (**Figure 5B**). Incubation with neutrophils led to significant decreases in the viability of GAS (**Figure 5A**) and *S. aureus* (**Figure 5B**), and the killing of GAS was significantly increased by the addition of pentamidine. This suggests the potential for pentamidine for sensitizing GAS to killing by the innate immune system.

**FIG 5.**
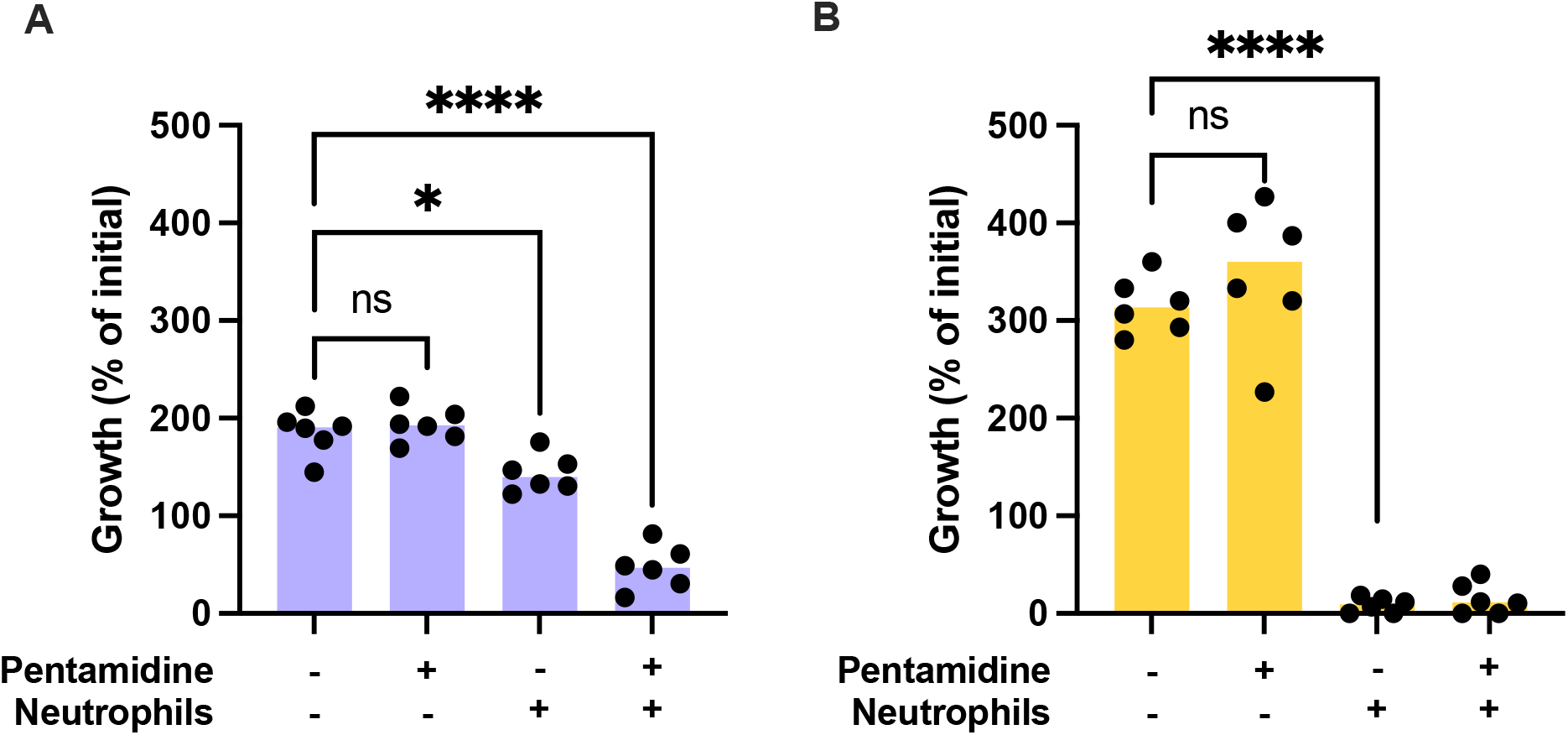
Pentamidine sensitized GAS to killing by neutrophils. Colony count of GAS (A) or *S. aureus* (B) after 2 hr infection of human primary neutrophils with 10 μM Pentamidine treatment. *P<0.05; ****P<0.00005 (ANOVA; Tukey post-test). Experiments were performed three times; error bars represent standard deviations.

## DISCUSSION

The Pathogen Box drug library has previously identified new inhibitors against a diverse set of pathogens including *Toxoplasma gondii* (55), *Candida albicans* (56), *Giardia lamblia, Cryptosporidium parvum* (57), *Vibrio cholerae* (58), *Acinetobacter baumannii* (59) and *E. coli* (60). In our study examining GAS, ∼2% of the compounds had significant protease inhibitor activity toward SpeB. A previous screen of 16,000 compounds of the Maybridge HitFinder HTS library had a hit rate of 0.018%, and identified a competitive inhibitor (52). Of our top 9 hits, activity against *Plasmodium falciparum* has been shown by MMV689758 and MMV676603 (61, 62), *Mycobacterium tuberculosis* by MMV687254 (63), *Candida auris, Trypanosoma evansi*, and *Schistosoma mansoni* by MMV022478 (64–66), and none could be found for MMV676588, MMV676512, MMV637953, MMV023953, and MMV024101. Altogether insufficient support could be found for any commonality in activity or mechanism between these compounds.

Pentamidine is a cationic aromatic diamine, which enters is known to interferes with polyamine synthesis and RNA polymerase activity in protozoal cells. It is typically delivered by oral, intravenous, and intramuscular routes, by topical delivery for skin disease and aerosolized delivery for pulmonary diseases have seen some success (67, 68). These results argue another potential utility for its use. Consistent with our findings, pentamidine has been observed to be an inhibitor of the gingipains of *Porphyromonas gingivalis* (69). These secreted virulence factors are, like streptopain, abundant and highly active cysteine proteases that target multiple host proteins to allow the bacterium to resist killing by neutrophils and other immune effector mechanisms (70).

Additional pathogens encode cysteine protease virulence factors, leaving the possibility that pentamidine may be more broadly useful as an inhibitor of bacterial virulence.

While other inhibitors of SpeB exist, such as the epoxide E-64 (N-[N-(L-3-trans-carboxyirane-2-carbonyl)-L-leucyl]-agmatine), their broad-spectrum activities can also inhibit essential host proteases (32). That pentamidine is already in clinical use argues that it can be tolerated. It has additionally been seen to prevent IL-1 cleavage, suggesting it can inhibit the host cysteine protease caspase-1 (71), without other obvious effects on the inflammasome (72). However, if this does occur in physiologically relevant conditions, it is likely to still not impair the immune responses because GAS is known to activate IL-1β and related cytokines independent of caspase-1 and other inflammasome-associated proteases (27, 32), and in some infection modes may even be protective (26). Lastly, there is significant overlap in the burden of the tropical diseases for which pentamidine is used and the burden of GAS. The long duration of pentamidine administration as a therapeutic or prophylactic typically required for these diseases has the possibility to impact whether an individual is co-infected by GAS. No existent clinical data could be found to support a correlation, which can be addressed in further studies examining disease incidence in these populations.

## ACKNOWLEDGEMENTS

We thank the Medicines for Malaria Venture foundation (MMV; Switzerland) for providing the Pathogen Box compounds that allowed this study. We appreciate the technical support provided by the Children’s Healthcare of Atlanta and Emory University’s Children’s Clinical and Translational Discovery Core for whole blood and cell processing. Molecular graphics and analyses performed with UCSF ChimeraX, developed by the Resource for Biocomputing, Visualization, and Informatics at the University of California, San Francisco, with support from NIH R01-GM129325 and the Office of Cyber Infrastructure and Computational Biology, NIAID. C.N.L. is supported by National Institutes of Health grants R01-AI153071 and R01-AI180089. C.N.L. is a Burroughs Wellcome Fund Investigator in the Pathogenesis of Infectious Disease. The content of this publication is solely the responsibility of the authors and does not necessarily represent the official views of any of its funders. No funders contributed to the study design or conclusions.

## Author contributions

K.T. and C.L., conducted the studies, wrote the manuscript, and approved the final manuscript.

